# Single-cell multiome regression models identify functional and disease-associated enhancers and enable chromatin potential analysis

**DOI:** 10.1101/2023.06.13.544851

**Authors:** Sneha Mitra, Rohan Malik, Wilfred Wong, Afsana Rahman, Alexander J. Hartemink, Yuri Pritykin, Kushal K. Dey, Christina S. Leslie

## Abstract

We present a novel gene-level regulatory model called SCARlink that predicts single-cell gene expression from single-cell chromatin accessibility within and flanking (+/-250kb) the genic loci by training on multiome (scRNA-seq and scATAC-seq co-assay) sequencing data. The approach uses regularized Poisson regression on tile-level accessibility data to jointly model all regulatory effects at a gene locus, avoiding the limitations of pairwise gene-peak correlations and dependence on a peak atlas. SCARlink significantly outperformed existing gene scoring methods for imputing gene expression from chromatin accessibility across across high-coverage multiome data sets while giving comparable to improved performance on low-coverage data sets. Shapley value analysis on trained models identified cell-type-specific gene enhancers that are validated by promoter capture Hi-C and are 8x-35x enriched in fine-mapped eQTLs and 22x-35x enriched in fine-mapped GWAS variants across 83 UK Biobank traits. We further show that SCARlink-predicted and observed gene expression vectors provide a robust way to compute a chromatin potential vector field to enable developmental trajectory analysis.

## Main

Multiome single-cell sequencing of chromatin accessibility and gene expression – where both scATAC-seq and scRNA-seq are read out from the same individual cells – has paved the way for novel computational methods that attempt to link enhancers to genes^1,2^, infer gene regulatory networks^3-5^, and resolve developmental trajectories based on the concept of chromatin potential, which proposes that accessibility at a locus precedes gene expression during differentiation^1^. At the most elementary level, several approaches exploit joint measurements of ATAC and RNA in single cells to identify pairwise correlations between individual accessible regions – defined as peaks or domains of open chromatin (DORCs) – and gene expression levels for enhancer-gene linking^1,6^. For example, a recent approach uses Poisson regression to test for pairwise correlation between peak accessibility and gene expression while also modeling batch or cell-specific covariates, with the goal of linking non-coding genetic variants that reside in such peaks to target genes^2^. Meanwhile, standard scATAC-seq analysis methods use simple scoring schemes to transform the data into a scRNA-like readout, based on aggregating chromatin accessibility near a gene promoter or across a genic locus to obtain an imputed gene expression value, to enable joint embedding of independently collected scATAC-seq and scRNA-seq data or transfer of clusters between the two^7^.

Motivated by these ideas, we propose SCARlink (Single-cell ATAC+RNA linking), a new gene-level predictive model for single-cell multiome data that predicts the expression of a gene from the accessibility of its genomic context in single cells (**Fig. 1a**). Unlike pairwise correlation approaches, our model captures the fact that elements both within the genic locus (e.g. intronic enhancers) and distal elements in flanking regions (+/-250kb by default) jointly regulate expression of the gene. We train the model using regularized Poisson regression on tile-level data to facilitate integration with standard preprocessing pipelines like ArchR^6^ and to avoid summarizing data as a peak atlas, which may miss events in rarer cell types. The regression coefficients across the genomic context can then be interpreted as identifying locations of functional enhancers across the single-cell data set. Moreover, we can use Shapley values, a well-known feature attribution method, to identify cell-type-specific enhancers, i.e. genomic tiles that are important for predicting expression across cells from a given cluster or annotation. Below, we show that our model outperforms existing methods for predicting single-cell gene expression from accessibility and correctly identifies cell-type-specific enhancers as validated by promoter-capture Hi-C. We further show that the regulatory regions determined using Shapley values from our modeling enrich for fine-mapped non-coding GWAS and eQTL variants. Finally, we demonstrate that using gene-level models for a set of developmentally regulated genes yield a robust implementation of the chromatin potential trajectory inference method.

**Figure 1.**
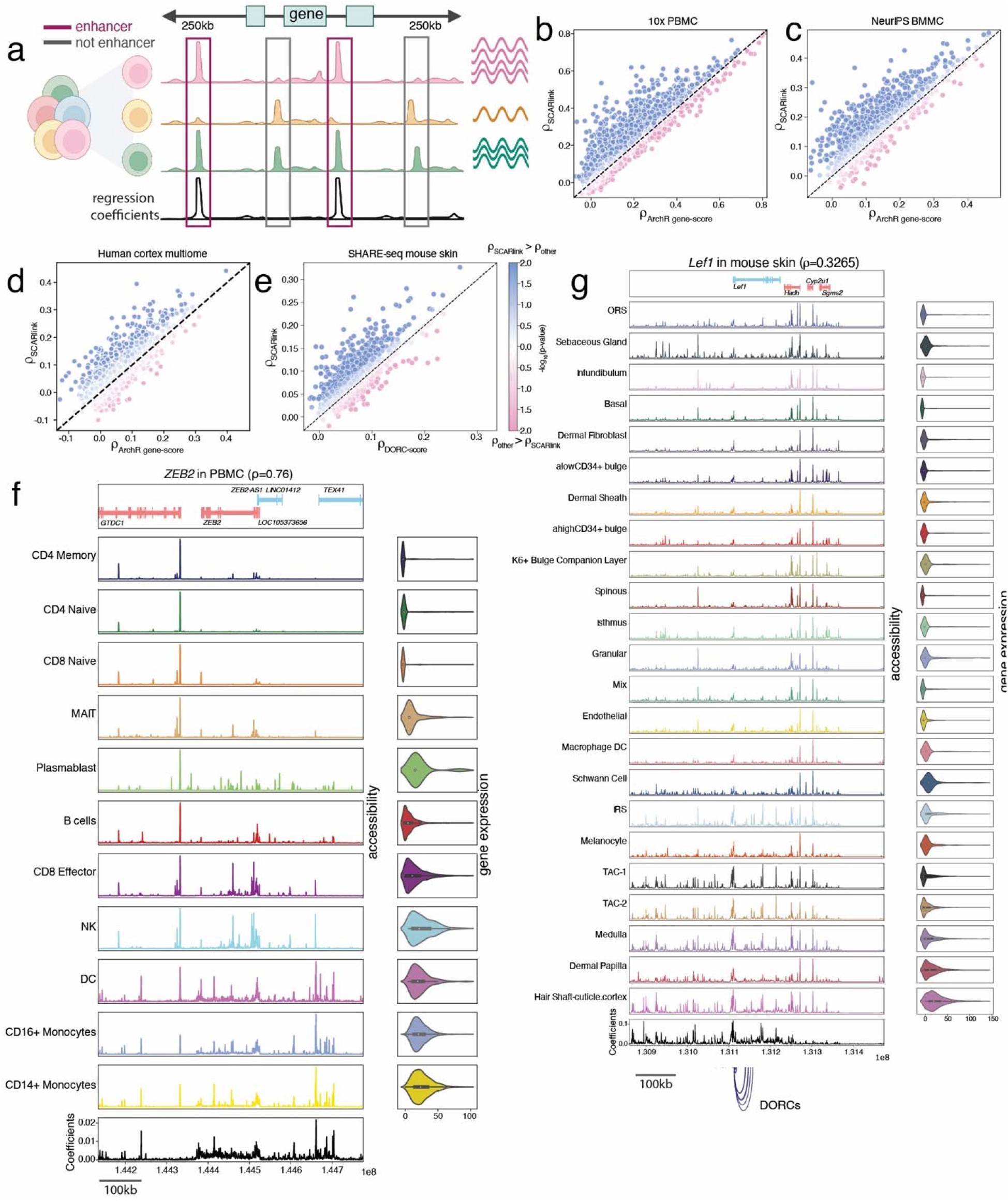
SCARlink accurately predicts single-cell gene expression from chromatin accessibility. **a**. The model takes as input single-cell ATAC-seq counts at a genic locus, aggregated over 500bp tiles spanning 250kb up/downstream and including the gene body, and uses regularized Poisson regression to predict the gene’s single-cell expression; both the scATAC-seq and scRNA-seq readouts are obtained from multiomic sequencing. The learned regression coefficients indicate the importance of each tile for predicting gene expression. **b-e**. Scatterplots showing Spearman correlation of predicted and observed gene expression for each gene using SCARlink vs. Spearman correlations using existing methods. Comparisons are performed against ArchR gene score predictions (**b-d**) on 10x PBMC, BMMC^8^, and developing human cortex^9^; and against DORC gene score predictions (**e**) on the mouse skin^1^ data set. **f**. Example model output for *ZEB2* from PBMC multiome data, showing regression coefficients at bottom and aggregated scATAC and scRNA by cell type. **g**. Example model output and comparison with annotated DORCs (shown using blue arcs below the coefficient panel) for *Lef1* from mouse skin SHARE-seq data.

SCARlink uses a regularized Poisson regression model on single cells to predict gene expression from chromatin accessibility. The chromatin accessibility is used as input in the form of 500bp tiles spanning a region from 250kb upstream to 250kb downstream of the gene body by default (**Fig 1a**). This genomic context is large enough to capture distal intergenic as well as intronic enhancers for most genes but can be extended or shortened as preferred. Since SCARlink is a gene-level model and genes are of variable length, the number of input tiles is different for every gene. We also constrain the model to learn positive regulatory elements by forcing the regression coefficients to be non-negative. While this is a limitation for identifying repressors, we found the regression coefficients to be more interpretable when we focused on enhancers.

We applied SCARlink to multiomic data sets of peripheral blood mononuclear cells (PBMC) from 10x Genomics, bone marrow mononuclear cells (BMMC)^8^, mouse skin^1^, developing human cortex^9^, pancreas^10,11^, and pituitary gland^12^. We ran the model on a subset of the top 5,000 most variable genes for each data set, filtered based on the sparsity of the gene expression vector (**Methods**). After filtering, we obtained 1,758 genes for PBMC, 786 genes for BMMC, 1,040 genes for mouse skin, 1,202 genes for developing human cortex, 792 genes for pancreas, and 1,222 genes for pituitary (**Supplementary Table 1**). For each gene-level model, we used Spearman correlation to compare the predicted gene expression to observed gene expression on held-out cells. We compared SCARlink against other available methods to predict single-cell gene expression from chromatin accessibility. One such method is the ArchR gene score, which aggregates accessibility across the gene body and flanking regions using an exponentially decaying function to downweight accessibility farther away from the gene. SCARlink significantly outperformed the ArchR gene score across all high-coverage data sets based on correlation with ground truth on held-out cells (one-sided signed-rank test over genes, p < 5.1e-96 on PBMC, p < 3.3e-90 on BMMC, and p < 1.3e-58 on developing human cortex). We also found SCARlink produced significantly higher correlations for a large fraction of individual genes in higher coverage data sets (44.4% of genes in PBMC, 56.8% of genes in BMMC, and 22.6% of genes in the developing cortex, at FDR < 0.05) as assessed by pairwise significance of correlation (**Methods**) (**Fig. 1b-d**). For lower coverage data sets, SCARlink performed comparably to the ArchR gene score on mouse skin multiome and pituitary (one-sided signed-rank test not significant in either direction) while outperforming it on pancreas (p < 1.8e-18, one-sided signed-rank test), albeit with fewer genes showing significantly better correlation (**Supplementary Fig. 1a-c, Supplementary Table 1**). In the human cortex multiome data, SCARlink outperformed another method of gene score prediction called ChrAccR that aggregates the accessibility in peaks near the TSS (p < 1.3e-93, one-sided signed-rank test; **Supplementary Fig. 1d)**. Domain of open chromatin (DORC) scores are computed by aggregating accessibility in peaks lying within 50kb and 500kb of the TSS that individually correlate with gene expression^1^. We found that our model yields predictions that are more correlated with expression than DORC scores in mouse skin (p < 8.9e-61, signed-rank test; significantly better performance on 36.6% of genes) (**Fig. 1e**), potentially because SCARlink is modeling the impact of chromatin accessibility across all tiles at once.

As an example to study the linkage between chromatin accessibility and gene expression, we used SCARlink to model regulation of *ZEB2* in the PBMC data set (**Fig. 1f)**. The learned regression coefficients across all the tiles (**Fig. 1f**, bottom) describe the regulatory importance of chromatin accessibility over the whole genomic context for *ZEB2*. Note that while SCARlink does not use cell type or cluster annotations as input, knowledge of clusters can be used to generate pseudobulk visualizations and thus interpret the regression coefficients. We also analyzed *Lef1* from mouse skin multiomic SHARE-seq data and found distal regions where high regression coefficients indicate that accessibility is correlated with transcription but which are not annotated as DORCs (near chr3:130,900,000; **Fig. 1g**). This highlights the advantage of SCARlink in using accessibility across all tiles for prediction of gene expression.

The regression coefficients generated using SCARlink indicate the overall importance of the accessibility in each tile for predicting gene expression across the data set. To quantify the contribution of each tile in the window for every cell type, we computed standardized average Shapley values per cell type on training cells (see **Methods** for computation of approximate Shapley scores under the SCARlink model). This allowed us to identify tiles as putative regulatory regions for the modeled gene in a particular cell type. We observed that predicted regulatory elements are most enriched within or in close proximity to the gene body (∼25kb) and decrease in prevalence in distal regions (**Supplementary Fig. 1e**). Because active enhancers are known to physically interact with promoters to enable transcription^13^, we hypothesized that SCARlink-predicted regulatory regions would be enriched for 3D interactions with the promoter of the modeled gene. Promoter capture Hi-C (PCHi-C) is a chromosome conformation capture assay that identifies promoter-interacting genomic regions using a genome-wide promoter bait library. We therefore sought to validate SCARlink-predicted regulatory regions across a subset of PBMC cell types using available hematopoietic PCHi-C^14^. We identified PCHi-C interactions in relevant cell types using a generalized additive model (**Methods**) and compared them to SCARlink-identified regions in T cell subpopulations, monocytes, and B cells in the PBMC multiomic data. As one example, we compared our Shapley values to PCHi-C interactions for the gene *HLA-DQB1* (**Fig. 2a**). We found that PCHi-C interactions in distal tiles display higher Shapley values than non-interacting tiles, particularly for B cells, a cell type in which *HLA-DQB1* is highly expressed (**Fig. 2a**). We then compared the Shapley values of tiles with and without PCHi-C interactions for highly expressed genes in each cell type (**Methods**) and confirmed that Shapley values for interacting tiles are significantly higher than for non-interacting tiles (**Fig. 2b**).

**Figure 2.**
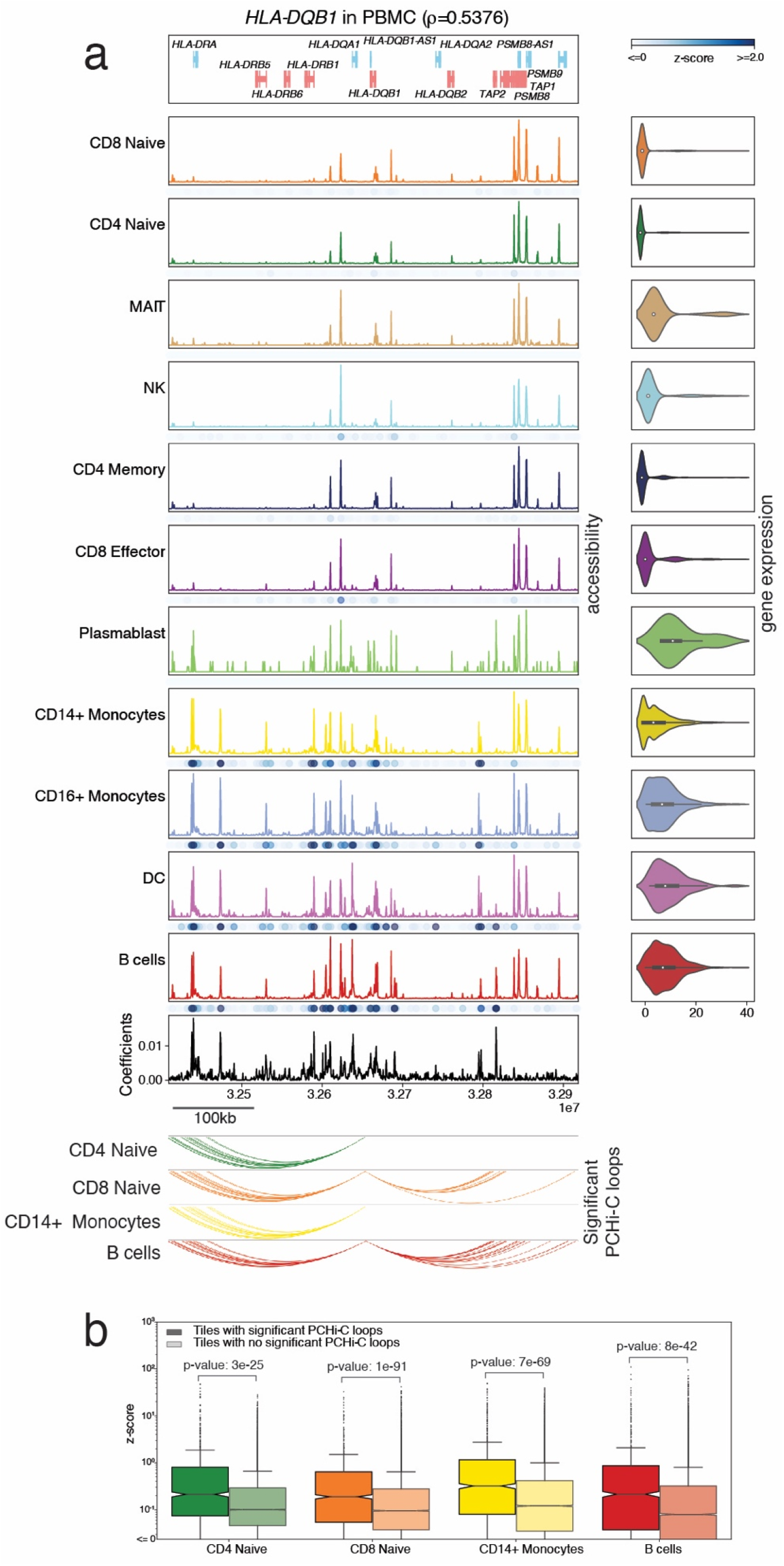
SCARlink coefficients enrich for promoter-linked chromatin interactions. **a**. SCARlink output of *HLA-DQB1* in PBMC multiome. Cell-type-specific standardized approximate Shapley scores (z-scores) of the tiles are plotted as blue dots under the accessibility panel of every cell type. Arc plots of significant PCHi-C interactions^14^ for *HLA-DQB1* of CD4 naïve T, CD8 naïve T, CD14+ monocytes, and B cells are shown below the model output. **b**. Boxplots comparing feature scores of tiles with or without PCHi-C interactions for highly expressed genes per cell type.

Next we assessed if the enhancer tiles predicted by SCARlink can be used to prioritize genetic variants causally associated with gene regulation and disease etiology. To this end, we first filtered a set of gene-linked tiles for each gene and cell type based on the significance of an approximate Shapley score (**Fig 3a, Methods**). We then performed an enrichment analysis of the resulting set of gene-linked tiles with respect to statistically fine-mapped eQTLs (PIP > 0.5) for the corresponding genes in the closest matched GTEx tissues^15^, and with respect to 10,164 statistically fine-mapped GWAS variants (PIP > 0.2) across 83 UK Biobank traits^16^ (average N=334,803) (**Supplementary Table 3**) in PBMC, pancreas, and pituitary gland (**Methods**). SCARlink gene-linked tiles in the three multiome datasets show 22x-35x enrichment of fine-mapped GWAS variants in the top 50,000 predicted gene-linked tiles (**Supplementary Table 4**) and outperformed a standard pairwise peak-gene linking implemented by ArchR. The enrichment increases with higher PIP thresholds (**Fig. 3b**). Moreover, the enrichment of the GWAS variants is individually higher for 64% of the 83 traits in SCARlink gene-linked tiles (**Supplementary Fig. 2a**). For the fine-mapped eQTL traits from matched GTEx tissues, we observed 15x-59x enrichment in PBMC for the first 15,000 gene-linked tiles (**Fig. 3c**, left) and 8x enrichment across predicted gene-linked tiles at FDR < 0.01 (**Fig. 3c**, right). We also observed 35x enrichment in pancreas multiome. Both PBMC and pancreas multiome gene-linked tiles have significantly higher enrichment than the enrichment using ArchR gene-linked peaks (**Fig. 3c**). To assess tissue-specific eQTL enrichment, we calculated the enrichment in PBMC and pituitary multiome of eQTLs from non-matching tissues from the GTEx database. We observed lower enrichment of eQTLs from other GTEx tissues (**Fig. 3d** and **Supplementary Fig. 2b**), suggesting that SCARlink is able to identify variants in regulatory regions that are tissue-specific and cell-type-specific.

**Figure 3.**
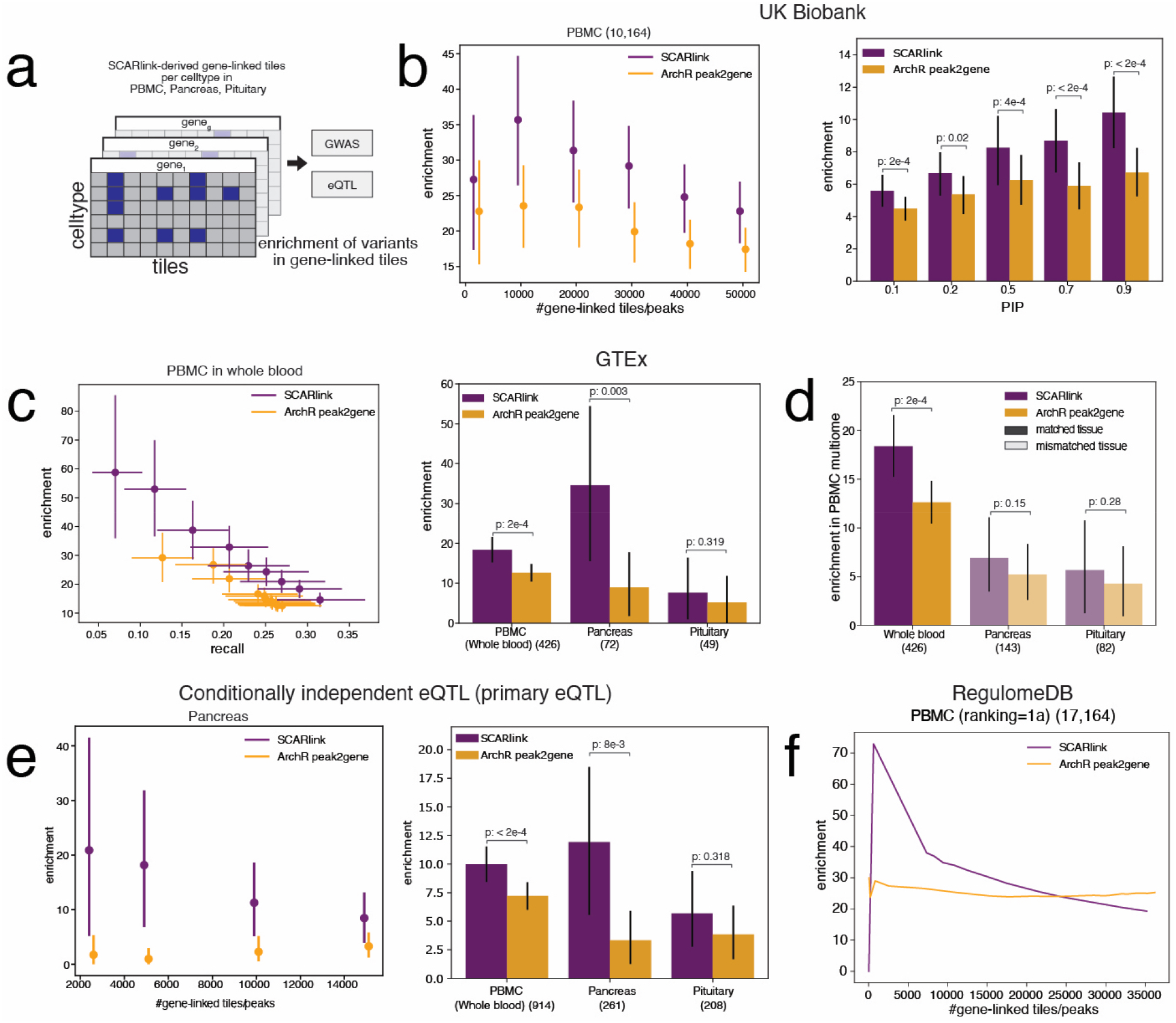
SCARlink-predicted gene-linked tiles enrich for causal variants. **a**. Schematic depicting the filtering of gene-linked tiles per cell type from SCARlink output of genes from PBMC, pancreas, and pituitary multiome. These filtered gene-linked tiles are then checked for enrichment of causal variants from GWAS, eQTLs, and other variant databases. **b**. Enrichment of 10,164 fine-mapped GWAS variants from UK Biobank in the gene-linked SCARlink tiles and ArchR peak2gene peaks as a function of the number of gene-linked tiles/peaks for PIP threshold of 0.2 (left). Comparison of enrichment at different PIP thresholds (right). The bars depicting 95% confidence interval were obtained by bootstrapping traits. **c**. Enrichment of 426 fine-mapped eQTLs from whole blood GTEx in PBMC multiome (left). Comparison of enrichment in the matched GTEx tissue as the multiome datasets (right). The number of fine-mapped variants per tissue is mentioned in brackets. **d**. Comparison of enrichment of eQTLs from GTEx tissues (pituitary, pancreas, and whole blood) in PBMC multiome. **e**. Enrichment of 261 primary independent eQTLs from pancreas as a function of number of gene-linked tiles/peaks (left). Enrichment of primary eQTLs in matched tissues in PBMC, pancreas, and pituitary (right). The bars depicting 95% confidence interval in **c-e** were obtained by bootstrapping genes. **f**. Enrichment of 17,164 variants from RegulomeDB of rank=1a in PBMC multiome.

We then assessed the enrichment of SCARlink gene-linked tiles in conditionally independent eQTL signals from GTEx. SCARlink showed 8x-21x enrichment of primary eQTLs (defined by the eQTL with the most significant association for the gene) in pancreas for the top 15,000 predicted gene-linked tiles and significantly higher enrichment in PBMC compared to ArchR peaks (**Fig. 3e)**. We additionally performed enrichment analysis of SCARlink gene-linked tiles with different categories of variants from RegulomeDB^17,18^. SCARlink showed higher enrichment for the top 25,000 gene-linked tiles in PBMC over ArchR peak-gene links for 17,164 RegulomeDB variants with a rank of 1a, corresponding to the most stringent cutoff based on motif accessibility at eQTL/caQTLs (**Fig. 3f**). SCARlink tiles also show higher enrichment for the top 10,000 gene-linked tiles in pituitary and top 5,000 tiles in pancreas (**Supplementary Fig. 2d-e, Supplementary Table 5**).

We also examined variants identified by statistical fine-mapping of eQTL and GWAS signals for which SCARlink provides evidence that the variant-containing tile is linked to the gene in specific cell types. This is to explore whether SCARlink can be used to identify putatively causal cell types for the variant action. One such variant, *rs112401631* (chr17:40608272:T:A), is a fine-mapped variant for asthma (PIP = 0.27) and is located in a tile that is significantly linked to the gene *CCR7* by SCARlink in various T cell subtypes (CD8 effector, CD4 memory, CD8 naïve, and CD4 naïve) (**Fig. 4a**). The *CCR7* gene is well known for its role in the homing of T cell populations to lymphoid organs^19,20^, and CCR7+ memory CD4+ T cells have previously been associated with severity of asthma^21,22^. A second example is the fine-mapped variant *rs12454712* (chr18:63178651:T:C) for insulin-like growth factor 1 (IGF1) and type 2 diabetes (adjusted by BMI) and lies in an intronic enhancer of *BCL2*^23^. IGF-1 is known to prevent apoptosis through the activity of *BCL2*, which encodes an anti-apoptotic transcription factor^24^. Furthermore, somatotropes secrete growth hormone that affects the production of IGF-1, and IGF-1 in turn negatively regulates growth hormone production^25^. Interestingly, we found this variant to be in a regulatory region of pituitary stem cells and somatotropes (**Fig. 4b**), possibly suggesting a role in pituitary stem cell differentiation. Additionally, both high and low IGF-1 levels have been associated with insulin resistance and a higher risk of type 2 diabetes^26^. While we found this variant within the regulatory region of cells from the pituitary gland, it is not accessible in the PBMC multiome (**Supplementary Fig. 3**), and SCARlink appropriately assigns the tile low significance in these cell types.

**Figure 4:**
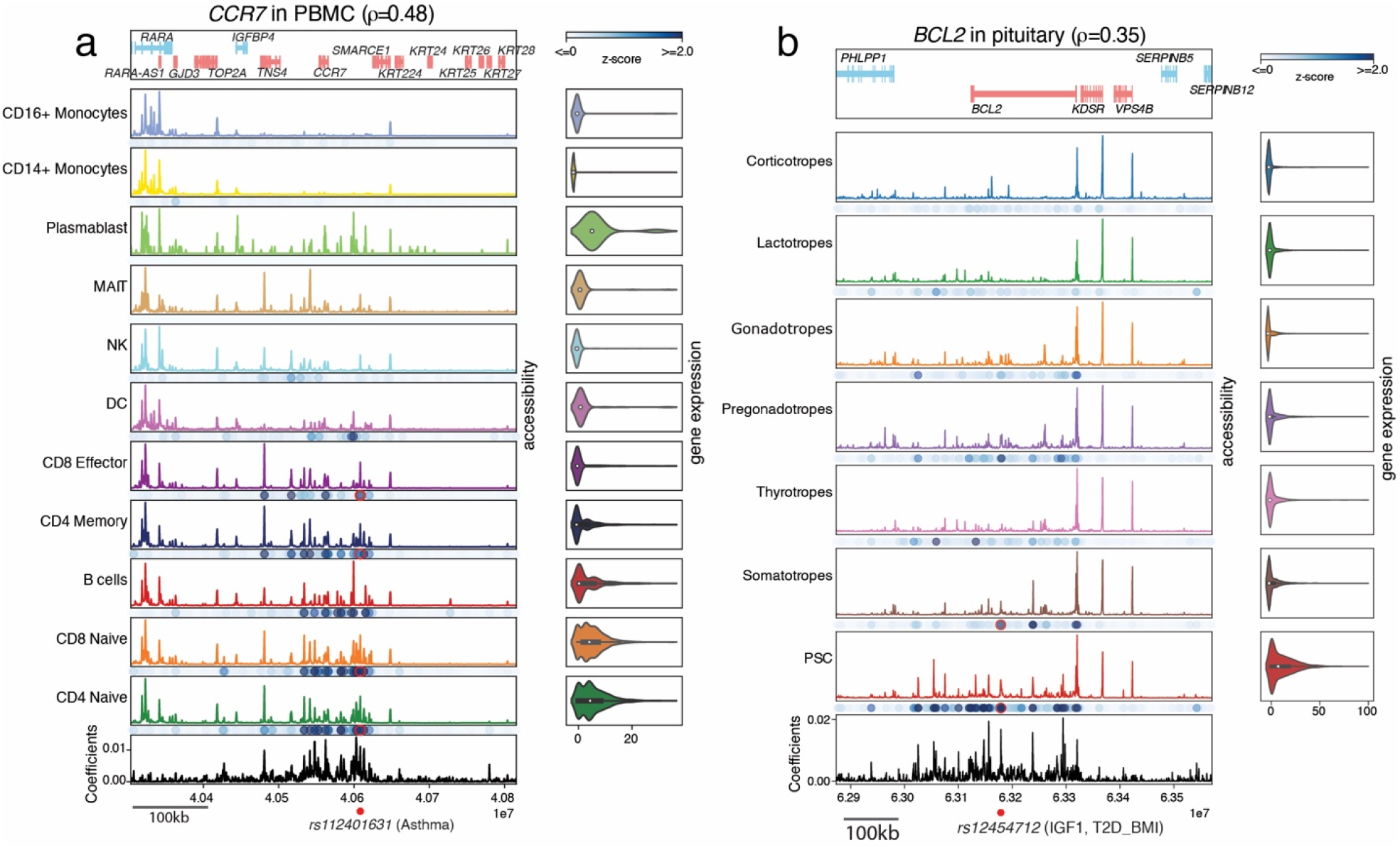
SCARlink-derived gene-linked tiles can reveal cell-type-specific disease-gene associations across tissues. **a**. SCARlink output of *CCR7* in PBMC. The red dot denotes a variant associated with asthma. The same position is highlighted in red under the cell types for which SCARlink predicted the variant-containing tile to be important. **b**. SCARlink output of *BCL2* in pituitary. The red dot at the bottom denotes variant associated with IGF1 and T2D_BMI. The tile containing the variant is highlighted in red for cell types for which SCARlink predicted the tile to be important.

We next asked whether SCARlink-identified regulatory regions become accessible before transcription of the modeled genes in developmental settings and thus can be used to determine the developmental trajectory through chromatin potential^1,9^. Analogous to the original definition of chromatin potential-based correlation between DORCs and genes, we computed a smoothed SCARlink-predicted gene expression vector for each given ‘source’ cell, identified a set of ‘target’ cells whose smoothed observed gene expression vectors are most correlated with the predicted source cell expression vector, and determined the corresponding chromatin potential vector from the source cell towards the average position of the target cells, and visualized in an FDL or UMAP embedding (**Methods**). We applied SCARlink in this fashion to derive chromatin potential vector fields for mouse skin, BMMC, pituitary gland, and developing human cortex. When computing chromatin potential, by default we chose all genes for which SCARlink-predicted gene expression was positively correlated with observed gene expression. This filtered out less than 5% of genes for mouse skin, BMMC, and pituitary, and 15% of genes from developing human cortex.

We found that the SCARlink chromatin potential vector fields recapitulate known differentiation trajectories in mouse skin, BMMC, and pituitary gland (**Fig. 5a-c**). However, in developing human cortex cells, chromatin potential failed to identify that the radial glia cell population is the root cell type^9^ (**Fig. 5d**). Upon comparing the difference between predicted and observed gene expression averaged over all genes, we found that this difference is the highest in the middle of the known developmental trajectory (nIPC/GluN1) and decreases afterwards (**Fig. 5d-g**). Examining further, we identified two clusters of genes based on hierarchical clustering of single-cell expression patterns (**Supplementary Fig. 4a, Supplementary Table 6**), with one cluster enriched for gene ontology terms related to glial cell differentiation (**Supplementary Fig. 4b-c**). Performing SCARlink chromatin potential analysis on this subset of 470 genes recovered the correct developmental trajectory (**Fig. 5h)**. For this subset of genes, we also found that the difference between average predicted and observed gene expression increases over the course of the trajectory, consistent with the opening of chromatin at these loci preceding target gene expression (**Fig. 5i-k**). While our analysis demonstrates the utility of chromatin potential as a strategy to identify a differentiation trajectory in multiome data sets, we also caution that prior selection of a subset of genes may be required to obtain results consistent with known biology.

**Figure 5.**
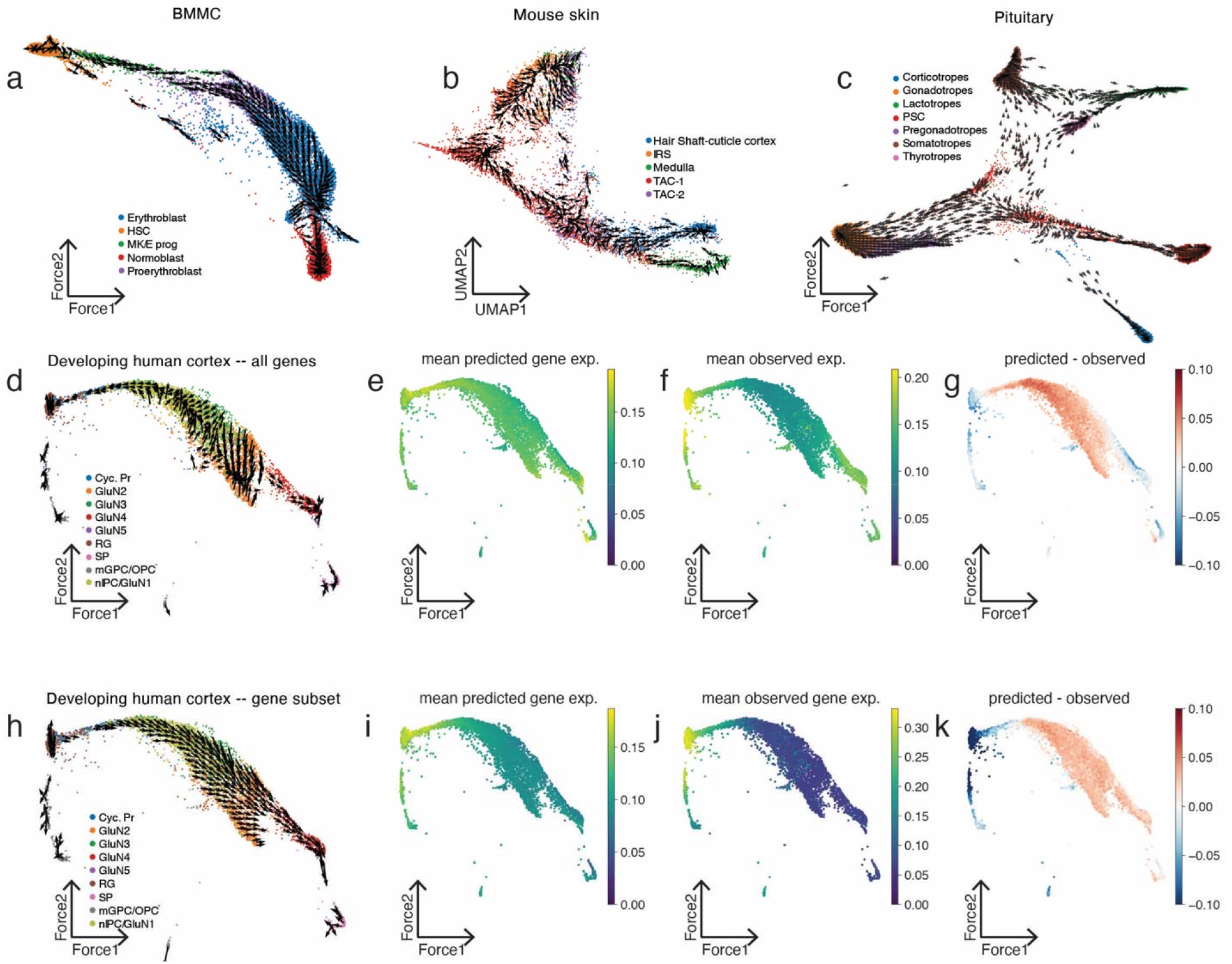
SCARlink provides a robust implementation of chromatin potential. **a-c**. SCARlink-computed chromatin potential applied to BMMC^8^, mouse skin^1^, and pituitary gland^51^ recapitulates known differentiation trajectory in each system. **d**. Chromatin potential does not capture the known differentiation trajectory of developing human cortex^9^ when using all genes with correlated predicted and observed gene expression. For the genes used in (**d**), **e-g** show the mean predicted expression, the mean observed expression, and the difference between the mean predicted and observed expression respectively. **h**. The known trajectory of the developing human cortex is better represented when only using a subset of the genes. For the genes used in (**h**), **i-k** show the mean predicted expression, the mean observed expression, and the difference between the mean predicted and observed expression respectively.

We have shown that SCARlink provides an effective and robust method for identifying cell-type-specific enhancers of genes without prior computation of a peak set. SCARlink also efficiently resolves the cell type specificity of tissue-relevant eQTLs and GWAS traits using Shapley value analysis and computes chromatin potential vector fields tracking development or differentiation. We note that SCARlink is designed to be a simple gene-level model, namely a (regularized) generalized linear model with a log link function and constrained to have non-negative regression coefficients. This simplicity enables fast training and model selection as well as very efficient computation of approximate Shapley values to identify significant tiles in a cell-type-specific manner. Additionally, by modeling additive positive effects, we obtain a highly interpretable model where significant tiles from Shapley analysis are validated by chromosome conformation capture data and enrich for fine-mapped eQTLs and GWAS variants. We also expect that SCARlink’s cell-type-specific enhancers and enhancer-gene links could be incorporated into functionally driven TWAS methods for predicting gene expression from genotype^27-30^. Despite the effectiveness of SCARlink’s generalized linear modeling, we can anticipate settings where more complex gene-level models might be suitable; for example, one could include interaction terms between tiles in the regression model or even employ non-linear neural network architectures for the same single-cell gene expression prediction task. Our implementation of SCARlink in TensorFlow should facilitate implementation of and comparison to these more complex models. Finally, there has been extensive work on DNA sequence models for bulk epigenomic and scATAC-seq data^31,32^, including in the context of prediction of bulk gene expression^33,34^. In future work, we plan to integrate DNA sequence information into SCARlink, sharing the sequence model associated with each cell across gene models, with the goal of modeling the regulatory grammar in enhancers as well as their regulatory impact on target gene expression.

## Methods

### Data preprocessing

Single-cell multiomic data was processed using Seurat^35^ (scRNA-seq) and ArchR^6^ (scATAC-seq). We performed quality control separately for scRNA-seq and scATAC-seq. We filtered out cells with mitochondrial reads > 20% for scRNA-seq with unannotated cell types (10x PBMC and pancreas). For scATAC-seq, we filtered for cells with at least 1,000 fragments and performed doublet detection on unannotated datasets. We performed CPM normalization of the scRNA-seq data. Then we ordered the cells in the same manner for both scRNA-seq and scATAC-seq. We selected the top 5,000 highly variable genes, using Seurat, and used this gene set as input to SCARlink.

### Cell type annotation

Cell type annotation was provided by the original studies for BMMC, developing human cortex, mouse skin, and pituitary gland. We performed cell type annotation of PBMC and pancreas using marker genes^35-37^.

### Gene regression model

SCARlink uses regularized Poisson regression to predict single-cell gene expression from single-cell chromatin accessibility.

We used ArchR to split the genome into 500bp tiles and computed tile-level scATAC-seq feature accessibility. We selected tiles that span 250kb up/downstream of and across the gene body. The accessibility within the tiles was normalized by the ReadsInTSS parameter, which is also the default normalization in ArchR, to control for sequencing depth and sample quality^6^. Gene expression values were normalized by counts per million (CPM). For each gene, the chromatin accessibility input to SCARlink was ReadsInTSS-normalized then min-max scaled on a per-tile basis across all cells. We ran the model separately on the 5,000 most variable genes determined using Seurat. Additionally, we filtered out genes for which the expression was too sparse with a threshold of 0.9, or 90% zeros.

We used L2 regularization with Poisson regression; i.e. for every gene, we optimized the following loss function:

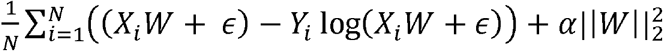

Here *N* corresponds to the number of cells, *X* corresponds to the min-max scaled accessibility matrix, Y corresponds to the gene expression vector, *W* is the learned regression coefficient vector, and *α* is the regularization parameter. We left out one-fifth of the data for testing. The regularization parameter was selected using 5-fold cross-validation on the remaining four-fifths of the cells. Spearman correlation was computed on the held-out test-cells. We used TensorFlow in Python to develop the model and the Adam optimizer for training. We constrained the regression coefficients to be non-negative, thereby learning only positive regulators for genes.

### Significance test for model predictions on individual genes

To compare overall performance of SCARlink predictions on test cells with other methods based on Spearman correlation with ground truth, we used a Wilcoxon signed-rank test over genes.

We also estimated whether the Spearman correlations of SCARlink predictions are significantly different from the correlations using other methods for individual genes. The correlations from the two methods are not independent because they are calculated on the same observed gene expression values. We calculated the following test statistic for each gene and performed a t-test to estimate significance^38^:

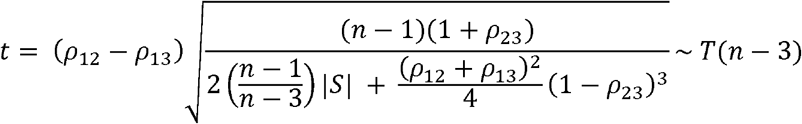

where

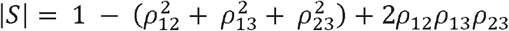

ρ_12_: Spearman correlation between SCARlink prediction and observed gene expression

ρ_13_: Spearman correlation between ArchR gene score/DORC score prediction and observed gene expression

ρ_23_: Spearman correlation between SCARlink prediction and ArchR gene score/DORC score prediction

n: number of cells in held-out test set

We performed FDR-correction of the p-values using the Benjamini-Hochberg method^39^. The scatter plots in **Fig. 1b-e** and **Supplementary Fig. 1a-c** are colored using these FDR-corrected p-values.

### Shapley scores and tile significance

After training the model, we used the SHAP Python package^40^ to compute Shapley values for a linear model, which closely approximate the Shapley values of our Poisson regression model:

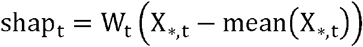

Here shap_t_ corresponds to the Shapley value of a particular tile_t_.

We computed these approximate Shapley values in a cell-type-specific manner. For each cell type, we iteratively sampled 50 training cells from the cell type to form a pseudobulk sample and computed Shapley values for each tile of the pseudobulk profile. We iterated 500 times and then averaged the Shapley values for each tile over iterations. This gave an averaged Shapley score for each tile and cell type. Finally, we standardized the scores using z-score transformation. We scaled features this way separately for each gene model in order to identify gene-linked tiles. Note that we estimate Shapley values only for cell types having at least 100 cells.

### PCHi-C analysis

We used publicly available PCHi-C data for hematopoietic cells^14^. We transformed the coordinates from hg19 to hg38 with liftOver^41^. Promoter Capture Hi-C loops at each promoter bait were identified by fitting a negative binomial generalized additive model^42^ to the observed counts as a function of GC content, mappability, and length of the restriction fragments alongside a smooth distance function parametrized using a reduced rank thin plate spline basis using the GAMLSS R package. If replicates were present, a replicate covariate was added to the model to control for library size. After this base model was fit, interactions were flagged by using the fitted distributions to compute a p-value. This overall strategy is akin to the GLM-based strategy of HiC-DC+ to identify significant interactions^43^. After p-values were computed for each restriction fragment in the vicinity of a promoter bait, p-values across replicates were pooled using Fisher’s method and corrected using Benjamini-Hochberg for each promoter bait. To further improve our ability to detect interactions, we employed locally adaptive weighting and screening to smooth the p-values and simultaneously control for the false discovery rate^44^.

For the Shapley value comparison, we used the *AverageExpression* function from Seurat^35^ to calculate average scaled gene expression and selected highly expressed genes per cell type. For every cell type, we restricted to genes with an average scaled gene expression of more than 0. Then we chose the top 50 genes if there were more than 50 highly expressed genes per cell type. Next we extracted all tiles that contain significant PCHi-C interactions for CD4 naïve T, CD8 naïve T, CD8 memory T, and B cells for these genes. If there were multiple tiles spanning one PCHi-C interaction, we selected the maximum Shapley value across the tiles. The background Shapley values are from tiles that do not contain any significant PCHi-C interactions for the same genes.

### Tile significance for variant analysis

We found the scaled Shapley scores were not comparable across gene models. Therefore, we used an additional metric to order the gene-linked tiles when computing enrichment-recall curves; specifically, we estimated the significance of difference in prediction of gene expression with and without a specific tile on held-out test cells using a paired Wilcoxon (signed-rank) test. We performed this significance test in a cell-type-specific manner across all genes in each multiome data set. The resulting p-values were then FDR-corrected using the Benjamini-Hochberg method^39^.

### ArchR peak2gene

We used ArchR^6^ to first perform peak calling using MACS2^45^ grouped by the cell type annotations. We then used the ArchR pipeline to link peaks to genes, which performs pairwise correlation of accessibility and gene expression on aggregated meta-cells. We used the same genomic window as SCARlink to predict the peak-gene links.

### GWAS enrichment analysis

We used fine-mapped GWAS variants from UK Biobank and first filtered out variants that lie within coding regions or are splicing eQTLs. UK Biobank originally has 94 traits. We retained the top 90% of the traits based on the number of fine-mapped variants lying within 250kb of all genes SCARlink was trained on. This resulted in 83 traits. We considered a variant to be a causal variant if it is associated with at least one trait with PIP > 0.2. This resulted in 10,164 variants that are present in tiles spanning 250kb upstream/downstream of all the genes from PBMC, pancreas, and pituitary. For each trait, we calculated precision as the ratio of the number of causal variants in predicted gene-linked tiles/peaks to the number of common variants in predicted gene-linked tiles/peaks. Then we calculated enrichment as previously described^2^, by dividing precision by the probability of encountering a causal variant of the given trait across all the tiles. We finally computed the average enrichment across all the traits:

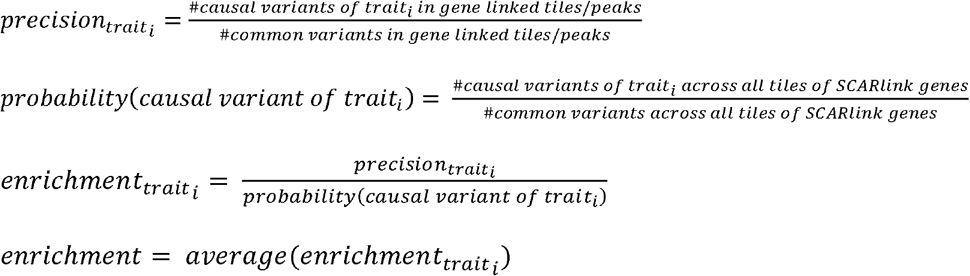

In the case of SCARlink gene-linked tiles, we restricted to genes having SCARlink-predicted gene expression correlation > 0.1 and to gene-linked tiles with p-value < 0.01. For ArchR gene-linked peaks, we restricted to peaks having correlation > 0.1 and FDR < 0.01.

### eQTL enrichment analysis

We used fine-mapped eQTLs from GTEx for whole blood, pancreas, and pituitary for computing enrichment in gene-linked tiles/peaks. We defined causal variants as having PIP > 0.5. Then separately for each gene and tissue, we computed precision, enrichment, and recall. We further computed the average enrichment and recall over genes per multiome data set:

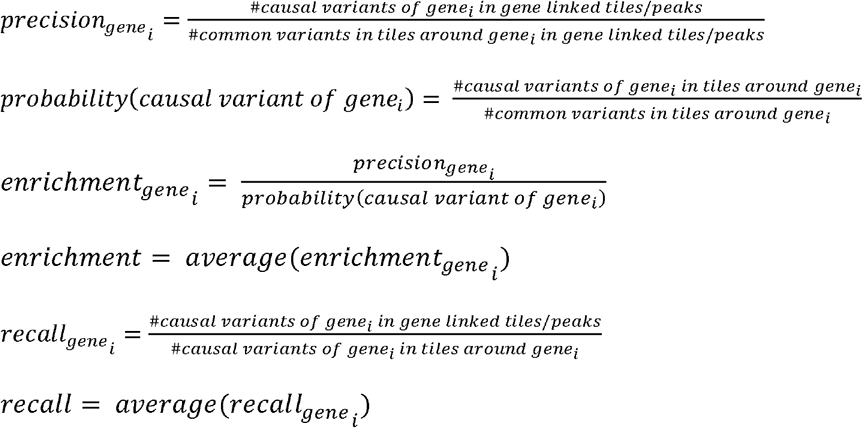

Additionally, we performed similar eQTL enrichment analysis on GTEx independent eQTLs for whole blood, pancreas, and pituitary. The primary independent eQTL is the most significantly associated variant^46^ and has a rank of 1. An eQTL with any other rank is an independent eQTL less important than the eQTLs with better ranks. There are at most 13 independent eQTLs, and the whole blood sample has more non-primary independent eQTLs than other tissues. We fixed a correlation cutoff of 0.1 for both SCARlink genes and ArchR peak2gene links and FDR < 0.01.

### RegulomeDB enrichment analysis

The variants in RegulomeDB^17,18^ are assigned ranks based on their associated regulatory features. Each variant is also assigned a probability score based on a random forest model, where probability scores are correlated with the ranks. We chose the most stringent set of variants with a rank of 1a, corresponding to variants associated with eQTL/caQTL and TF binding with matched motif, footprint, and accessible chromatin. We further restricted to variants with a probability score > 0.9. We considered these variants to be the putative regulatory variants.

We computed enrichment as follows:

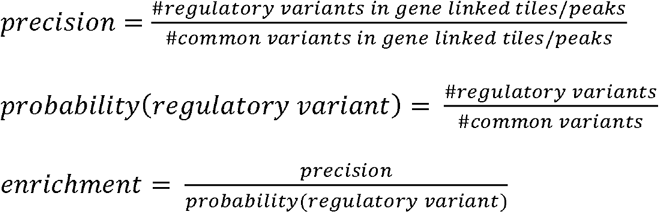

### Chromatin potential using SCARlink

We ran chromatin potential on smoothed SCARlink-predicted and observed gene expression values. Smoothing was performed over a k-nearest neighbor (kNN) graph (k=50) built using a lower dimensional representation of the scATAC-seq data based on latent sematic indexing (LSI) from ArchR. We retained the genes for which the predicted and observed gene expression are positively correlated. We then scaled the smoothed predicted and observed gene expression using min-max scaling. Following this, as in the published chromatin potential approach^1^, for each cell *i* in the predicted space, we identified the nearest neighbors (k=10) in the observed space:

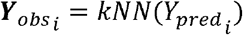

Here, 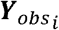 is the scaled and smoothed observed expression matrix of the 10 cells with the highest correlation with the scaled and smoothed predicted expression vector of cell *i*, 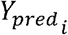. We then plotted chromatin potential arrows on the force directed layout (FDL) or UMAP from each cell *i*, to the average position of the cells corresponding to 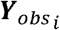. These arrows are further smoothed over a grid layout on the FDL/UMAP embedding.

We used FDL visualizations for all data sets except mouse skin, where we used the previously published UMAP ^1^. Additionally, for the mouse skin data, we ran the analysis on a subset of cell types to compare to reported results^1^.

By default, we do not filter out any genes except the ones with negative correlation between predicted and observed expression. We found that by using all genes, we could not always obtain the known differentiation trajectory, as in the case of developing human cortex. In this data set, we performed hierarchical clustering of genes based on cosine distance of observed gene expression vectors across all cell types, identified two clusters, and repeated chromatin potential analysis with genes in one of the clusters.

## Data availability

We downloaded the PBMC multiome from 10X Genomics. BMMC data was part of the NeurIPS 2021 open problem, and the data set was downloaded from GEO (GSE194122). We used BMMC samples labeled as site1_donor1, site1_donor2, site1_donor3, site2_donor1, site2_donor4, site2_donor5, site3_donor10, site3_donor6, site3_donor7, and site4_donor9. These samples showed the least batch effect. Mouse skin SHARE-seq data and DORC annotations were downloaded from GEO (GSE104203). The UMAP used for mouse skin was shared by the authors^1^. Pituitary multiome data was downloaded from GEO (GSE178454). The developing human cortex scRNA-seq was downloaded from GEO (GSE162170) and the corresponding multiomic scATAC-seq was downloaded from links listed in https://github.com/GreenleafLab/brainchromatin/blob/main/links.txt. We used samples labeled hft_ctx_w21_dc2r2_r1 and hft_ctx_w21_dc2r2_r2 with the least batch effect. We downloaded the pancreas multiome data set from the ENCODE portal (multiomic series ENCR316WAS).

We used common variants from the 1000 Genomes Project, phase 3^47^. The fine-mapped eQTLs for whole blood, pancreas, and pituitary were downloaded from GTEx v8^15^. The fine-mapping was performed using CAVIAR ^48^. We also downloaded the conditionally independent eQTL from GTEx v8. UK Biobank GWAS data with fine-mapping using SuSIE^49^ and FINEMAP^50^ was downloaded from the Finucane lab (https://www.finucanelab.org/data).

## Code availability

SCARlink is available on GitHub at: https://github.com/snehamitra/SCARlink/.

