## Supplementary Fig. for "Single-cell multiome regression models identify functional and disease-associated enhancers and enable chromatin potential analysis"

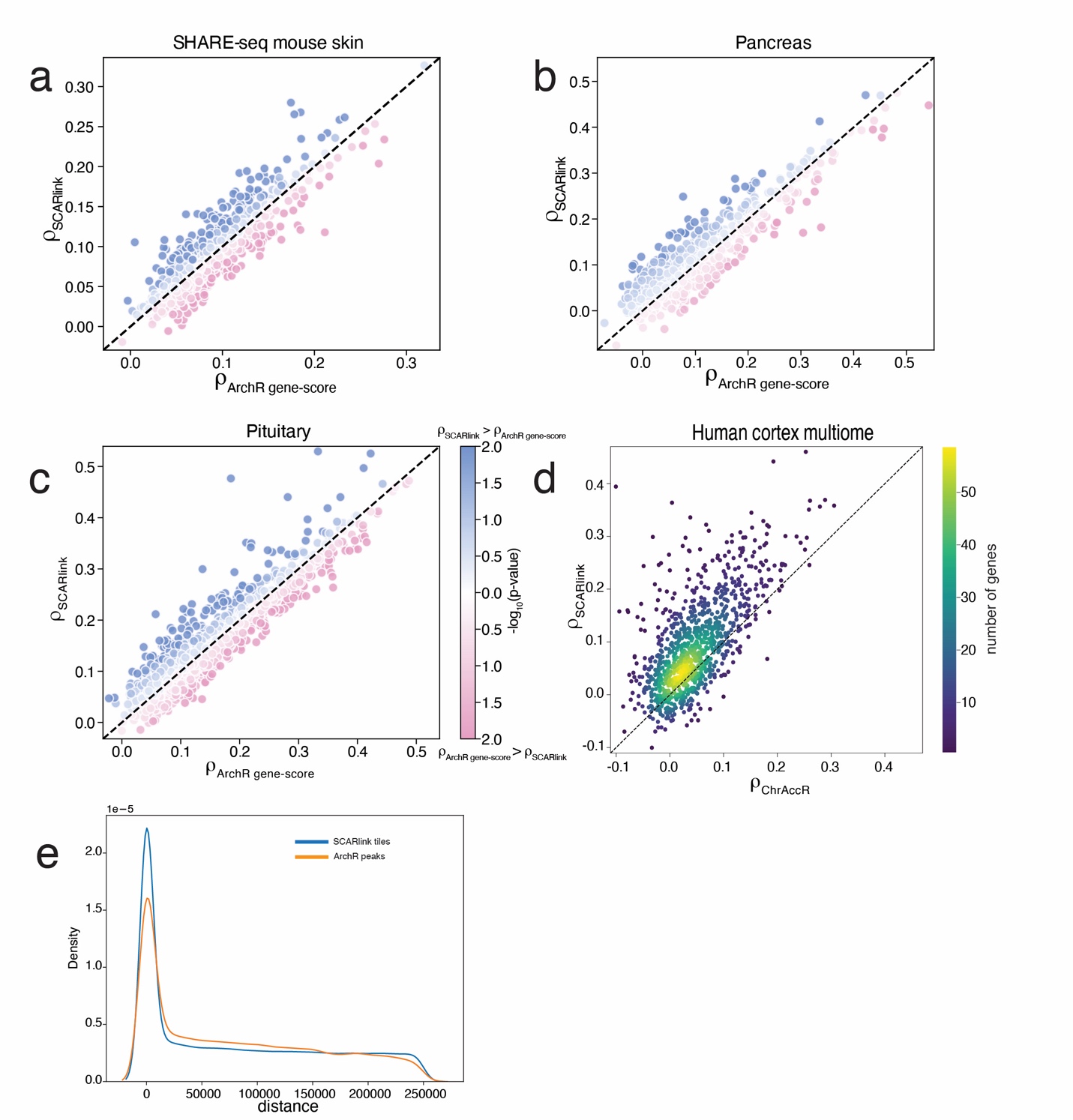

**Supplementary Figure 1**. **SCARlink prediction of gene expression compared to existing methods.** Comparison of Spearman correlation of prediction of gene expression between SCARlink and ArchR gene score in (**a**) mouse skin^1^, (**b**) pancreas^2,3^, and (**c**) pituitary multiome^4^. For each of the scatterplots in **a-c**, a significance score is computed between the Spearman correlations of SCARlink and ArchR gene score. The dots are colored based on the p-values. (**d**) Comparison of Spearman correlation of prediction of gene expression between SCARlink and ChrAccR in developing human cortex multiome. Each dot is a single gene. We use the Spearman correlations pre-computed using ChrAccR scores for each gene as previously reported^5^. Hence, no statistical tests were performed. So here the dots are colored based on the density of genes. **e.** Smoothed distribution plot depicting the number regulatory regions predicted within SCARlink tiles (blue) and ArchR paired peak-gene links (orange) is highest within the gene body (dist=0) and decreases on moving further away from the gene body.

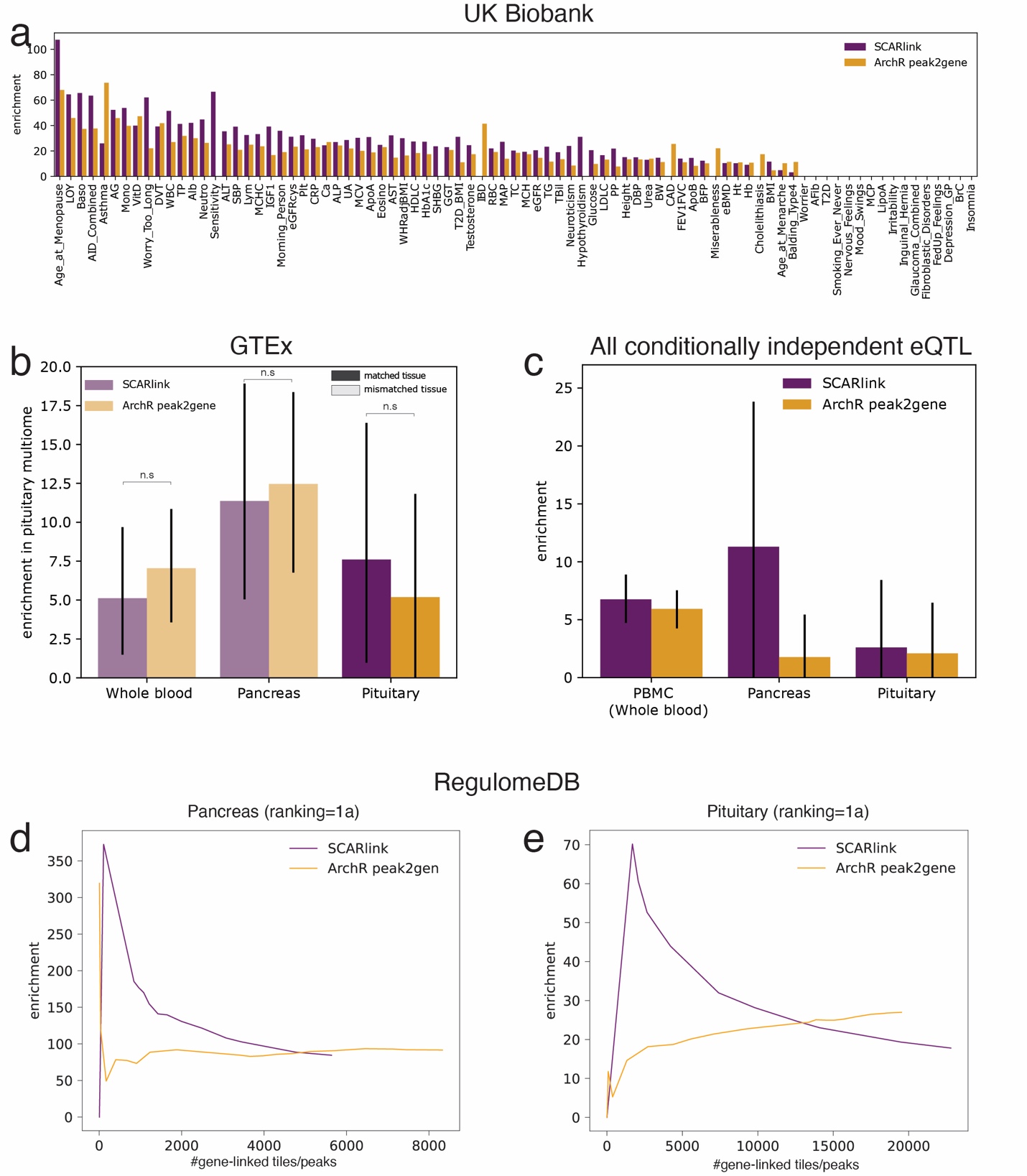

**Supplementary Figure 2.** **Comparison of variant enrichment in gene-linked tiles/peaks from SCARlink and ArchR peak2gene. a.** Trait specific enrichment of fine-mapped GWAS variants from UK Biobank for PIP > 0.2. Enrichment for a trait is 0 when no variants are found in predicted gene-linked tiles/peaks. **b.** Enrichment of pituitary multiome for GTEx pituitary, and other GTEx tissues (pancreas and whole blood). **c.** Enrichment plots for all independent eQTLs in PBMC, pancreas, and pituitary for closely linked tissues. Enrichment plots for RegulomeDB variants with ranking=1a in **d.** pancreas and **e.** pituitary multiome.

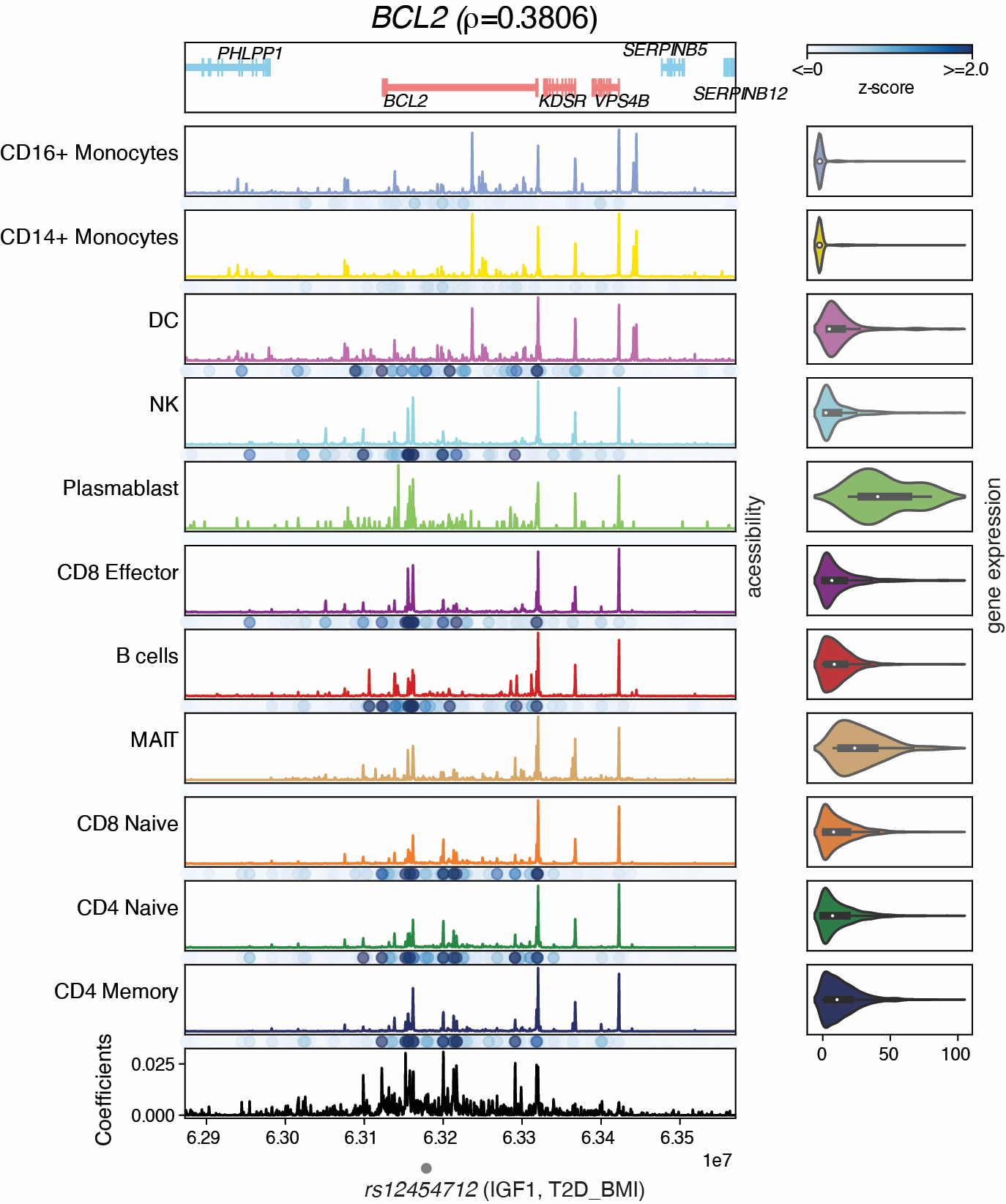

**Supplementary Figure 3. SCARlink output of BCL2 in 10x PBMC.** The grey dot at the bottom denotes the variant associated with IGF1 and T2D_BMI. The variant-containing tile is not important across any of the cell types in PBMC.

­­­­­­
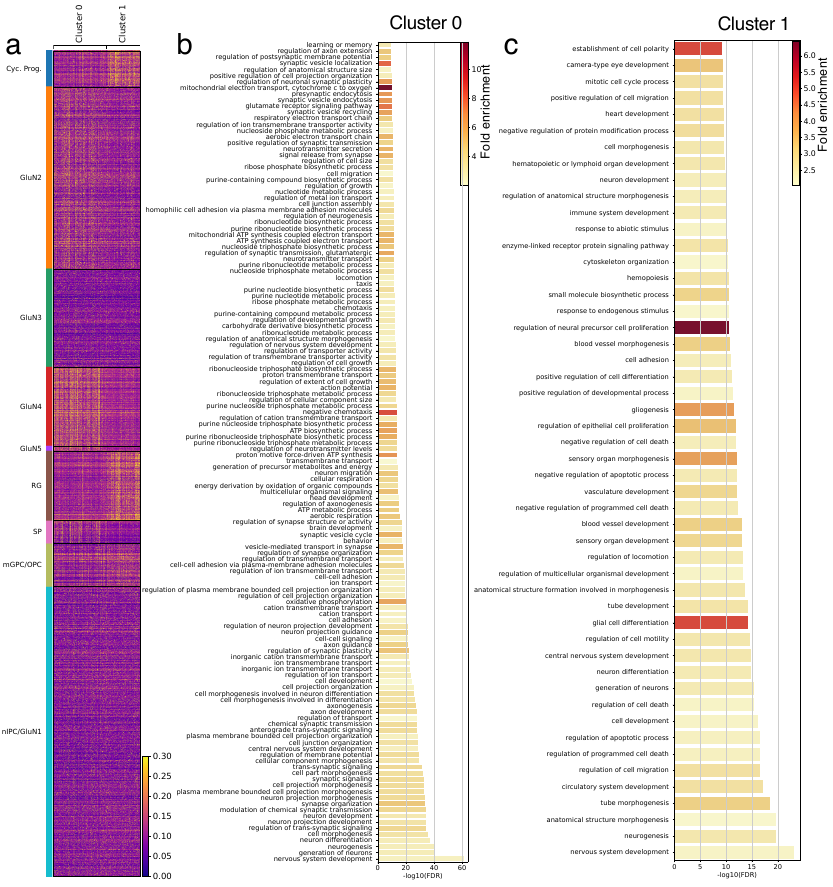

**Supplementary Figure 4. Enriched GO terms for the two clusters of genes in developing human cortex. a.** Heatmap showing comparison of gene expression in the two identified clusters across the cell-types. GO terms enriched in **b.** cluster 0 and **c.** cluster 1. Genes in cluster 1 capture the known differentiation trajectory.
